# Role of astroglial ACBP in energy metabolism flexibility and feeding responses to metabolic challenges in male mice

**DOI:** 10.1101/2022.09.01.506231

**Authors:** K. Bouyakdan, R. Manceau, J. Robb, D. Rodaros, S. Fulton, T. Alquier

## Abstract

Acyl-CoA Binding Protein (ACBP), also known as Diazepam Binding Inhibitor (DBI), has recently emerged as a hypothalamic and brainstem gliopeptide regulating energy balance. Previous work has shown that the ACBP-derived octadecaneuropeptide exerts strong anorectic action via POMC neuron activation and the melanocortin-4 receptor. Importantly, targeted ACBP loss-of-function in astrocytes promotes hyperphagia and diet-induced obesity while its overexpression in arcuate astrocytes reduces feeding and body weight. Despite this knowledge, the role of astroglial ACBP in adaptive feeding and metabolic responses to acute metabolic challenges has not been investigated. Using different paradigms, we found that ACBP deletion in Glial Fibrillary Acidic Protein (GFAP)-positive astrocytes does not affect diet-induced weight loss in obese male mice nor metabolic parameters in chow-fed mice (e.g. energy expenditure, body temperature) during fasting, cold exposure and at thermoneutrality. In contrast, astroglial ACBP deletion impairs meal pattern and feeding responses during refeeding after a fast and during cold exposure, thereby showing that ACBP is required to mount an appropriate feeding response in states of increased energy demand. These findings challenge the general view that astroglial ACBP exerts anorectic effects and suggest that regulation of feeding by ACBP is dependent on metabolic status.

## INTRODUCTION

Hypothalamic neurocircuits play a key role in the maintenance of energy homeostasis by regulating adaptive feeding and metabolic responses to changing energy needs (1). The melanocortin system in the arcuate nucleus (ARC) is critical in this control and is composed of two functionally opposed neuronal populations: agouti-related peptide (AgRP) neurons and proopiomelanocortin (POMC) neurons (2). POMC neurons are typically activated by metabolic signals during states of positive energy balance to reduce feeding and enhance energy expenditure via the release of alpha-Melanocyte Stimulating Hormone (alpha-MSH) and activation of the melanocortin-4 receptor (MC4R) (3). Accumulating evidence shows that hypothalamic astrocytes also respond to metabolic signals (4–8) and regulate parameters of energy balance in part through the hypothalamic melanocortin pathway (reviewed in (9, 10)). The mechanisms by which astrocytes regulate the activity of AgRP and POMC neurons involve changes in the astroglial coverage of neurons and synapses (8, 11, 12) and the secretion of metabolites (11, 13, 14). Recently, we and others have shown that astroglial acyl-CoA Binding Protein (ACBP), also known as Diazepam Binding Inhibitor (DBI), is a gliopeptide that regulates the activity of hypothalamic and brainstem neurocircuits controlling energy balance (15). ACBP plays a dual role in astrocytes: as an intracellular protein regulating long chain Acyl-CoA metabolism (16, 17) and as a secreted peptide acting on neurons (18). Expression of ACBP in the hypothalamus is upregulated by signals of energy sufficiency while decreased by fasting in a circadian manner (19–21). Upon release, ACBP is cleaved to produce endozepines including the octadecaneuropeptide (ODN) which signals through an unidentified ODN GPCR to reduce food intake (22). Central administration of ODN has a strong anorectic action in goldfish and rodents that is dependent on POMC neuron activation and MC4R in the ARC and brainstem (15). ODN increases the activity of POMC neurons in the ARC and brainstem via the unidentified ODN GPCR (19, 23). In addition, central ODN acutely increases glucose tolerance in rats (20) and reverses diet-induced obesity when administered chronically (23). ACBP in tanycytes exerts similar effects in addition to promoting the transport of circulating leptin to the hypothalamic parenchyma (23). We recently showed that astroglial ACBP deletion throughout the brain heightens susceptibility to diet-induced obesity in male and female mice, a phenotype that was prevented by the rescue of ACBP specifically in ARC Glial Fibrillary Acidic Protein (GFAP) positive astrocytes (19). Despite this evidence supporting the role of ACBP and ODN in feeding and obesity, the role of astroglial ACBP in energy metabolism flexibility and adaptive feeding responses to acute metabolic challenges has not been investigated. Here we used multiple paradigms to test if endogenous ACBP regulates metabolic flexibility and feeding behavior in male mice with targeted deletion of ACBP in astrocytes.

## RESULTS-DISCUSSION

### Astroglial ACBP deletion does not affect diet-induced weight loss in obese mice

We previously reported that male and female ACBP^GFAP^ KO mice are more prone to diet-induced obesity (DIO) during a high-fat diet (HFD) regimen compared to ACBP^GFAP^ WT (GFAP^Cre^) control mice (19). We sought to determine if ACBP deficiency affects the metabolic adaptation responses to the transition to a normal chow diet in HFD obese mice. Replicating our previous findings, we found that body weight and food intake are increased in male ACBP^GFAP^ KO mice fed with the HFD for 14 weeks (Fig 1A), with hyperphagia preceding the onset of overweight (Fig 1B). A slight but non-significant reduction of energy expenditure was observed before mice became overweight (after 6 weeks of HFD) (Fig 1C) with no change in respiratory exchange ratio (RER) (Fig 1D). Together, these results suggest that increased susceptibility to DIO of ACBP^GFAP^ KO mice is mostly due to increased food intake. Obese control littermates and ACBP^GFAP^ KO male mice were then transitioned to a chow diet for 4 weeks. Within the first week of the diet transition, body weight and food intake decreased similarly in both groups and remained stable thereafter (Fig 1E and F). Together these findings suggest that the obesity-prone phenotype in ACBP^GFAP^ KO mice is reversible and that astroglial ACBP deficiency does not affect diet-induced weight loss in obese male mice.

**Figure 1:**
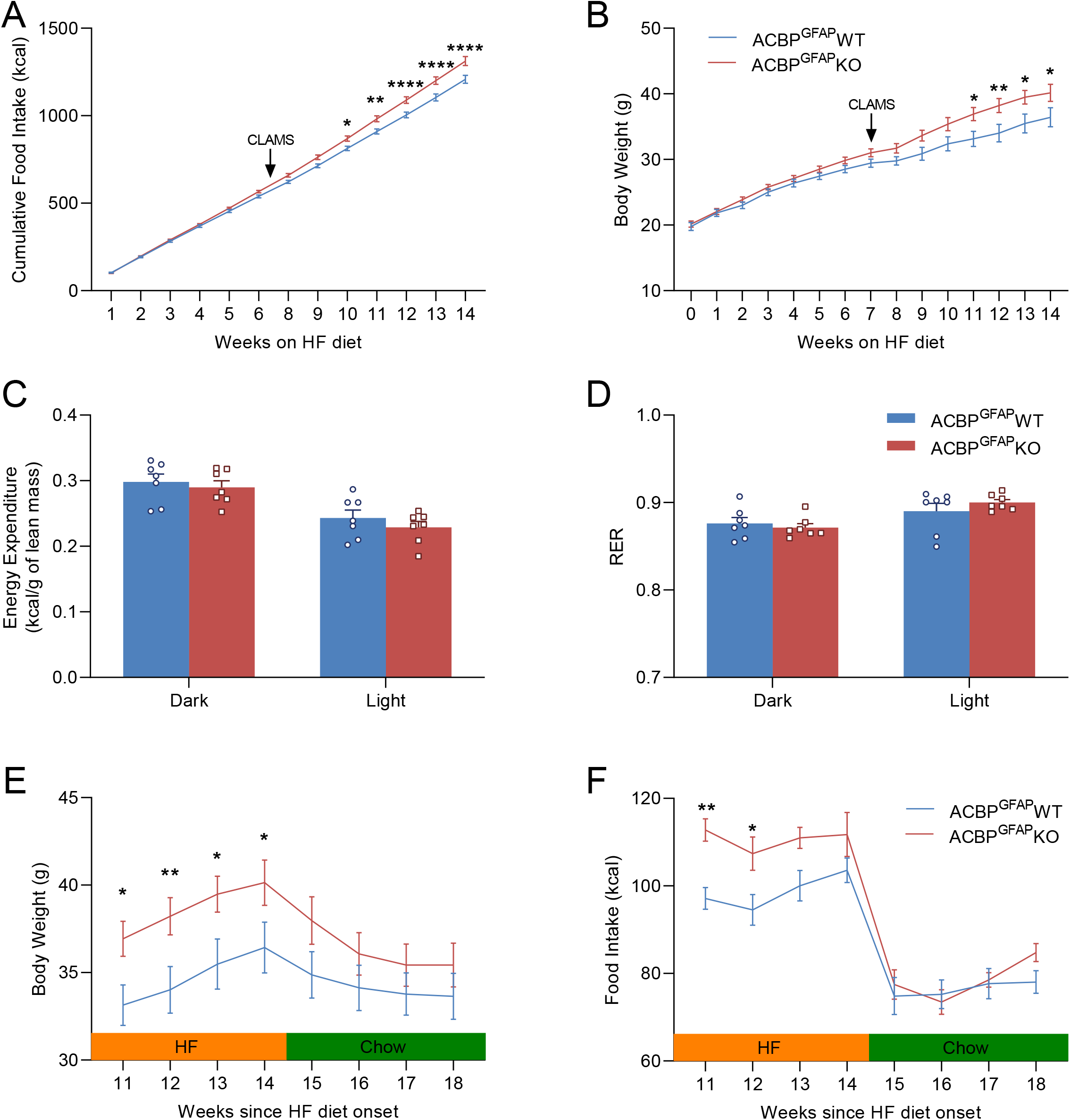
Astroglial ACBP deletion does not affect body weight loss in diet-induced obese mice. (**A**) Cumulative food intake and (**B**) body weight of ACBP^GFAP^ WT and KO male mice fed with a high fat diet (HFD) during 14 weeks. Mice were housed in metabolic cages during week 7 (indicated by the arrow). (**C**) Energy expenditure (normalized to lean mass) and (**D**) respiratory exchange ratio (RER) were measured in CLAMS metabolic cages during 48h following a 24h acclimation period. (**E**) Body weight and (**F**) food intake (kcal) are shown from week 11 to week 14 of the HFD regimen followed by a switch to regular chow diet for 4 weeks. * p<0.05, ** p<0.01, **** p<0.0001, Two-way ANOVA with Sidak post hoc test compared to ACBP^GFAP^ WT, N=7.

### Deletion of ACBP in astrocytes impairs feeding responses after a fast

We and others have shown that ACBP expression in the hypothalamus is regulated in a circadian manner and reduced by fasting (19, 21). In contrast, ACBP expression and secretion is increased by signals of positive energy balance including glucose and insulin (20, 23). In addition, central administration of the ACBP-derived peptide ODN regulates blood glucose levels and glucose tolerance in a manner dependent on the feeding status (20). Consistently, we observed increased whole body glucose utilization after intracerebroventricular (ICV) administration of ODN (19). Together, these findings suggest that astroglial ACBP regulate metabolic and feeding adaptive responses to a fast. To test this hypothesis, different cohorts of male ACBP^GFAP^ KO and control ACBP^GFAP^ WT mice were subjected to a 16h fast followed by *ad libitum* refeeding while housed in metabolic cages. Feeding behavior and meal pattern were analyzed in both *ad-libitum* fed and refed conditions. The total number of meals was not affected by feeding status or genotype (Fig 2A) while the average meal size (Fig 2B) and time spent eating (Fig 2C) were elevated during refeeding in both genotypes. Importantly, cumulative food intake was not greater in ACBP^GFAP^ KO mice during refeeding as compared to control mice (Fig 2D) thereby suggesting that ACBP is required to mount a normal feeding response after a fast. Based on our current (Fig 1B) and previous findings showing that ACBP and ODN exert an anorectic action (19, 24), an increased cumulative food intake would have been expected during refeeding in ACBP^GFAP^ KO mice.

**Figure 2:**
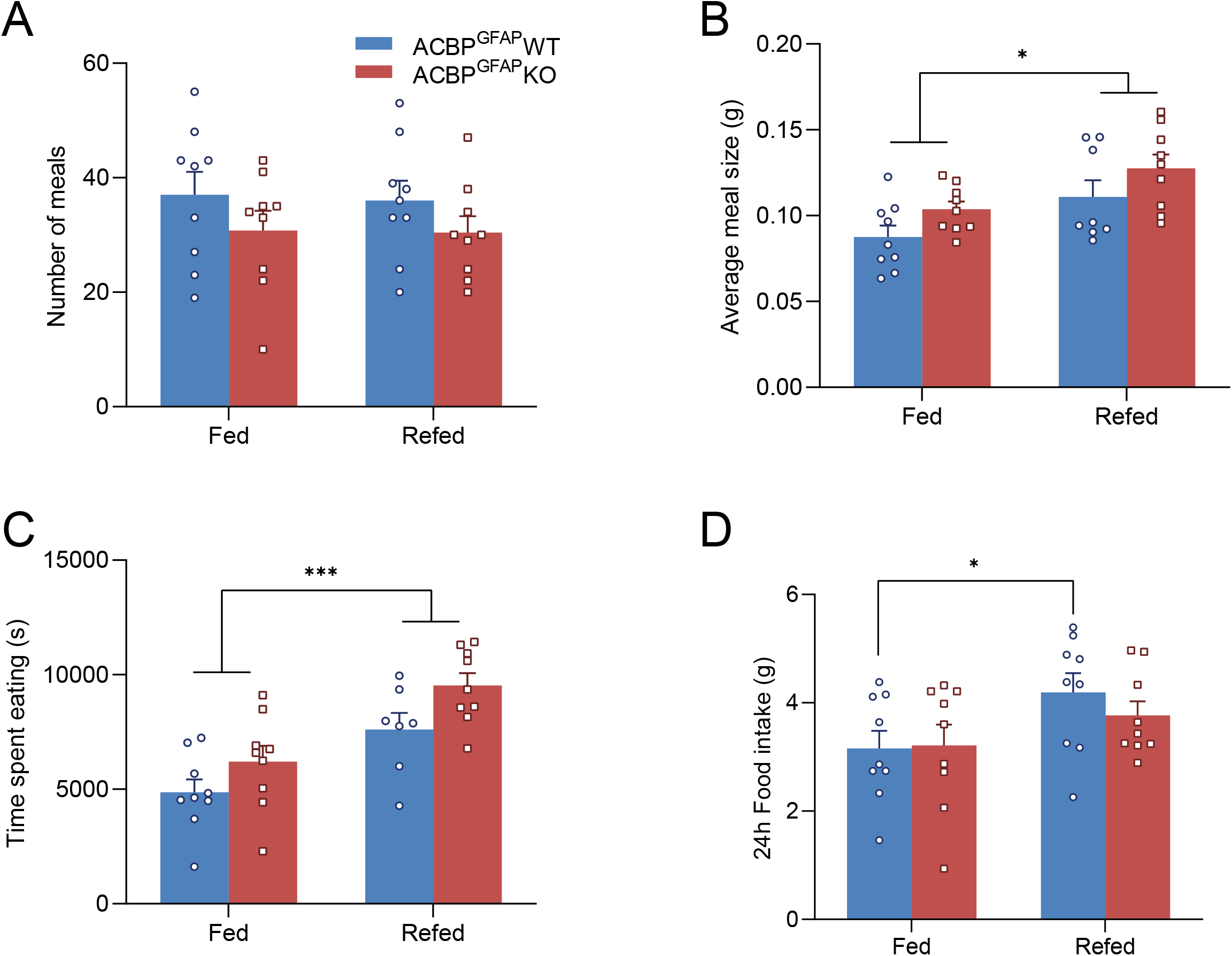
Meal pattern in astroglial ACBP knockout mice during refeeding after a fast. ACBP^GFAP^ WT and KO male mice were subjected to a 16h fast in CLAMS metabolic cages. (**A**) Total number of meals, (**B**) Average meal size, (**C**) Total time spent eating and (**D**) Cumulative Food intake were measured during the 24h before and the 24h following the fast. * p<0.05, *** p<0.001, Two-way ANOVA with Sidak post hoc test compared to fed state, N=9.

During the fasting-refeeding protocol, blood glucose was measured to assess potential changes in glucoregulatory responses in a different cohort. As expected, glycemia was blunted in the fasted state and elevated during the refeeding period in control mice (Fig 3A). Similar changes in blood glucose were observed in ACBP^GFAP^ KO mice to suggest that ACBP deletion does not affect the regulation of glycemia during a fast and subsequent refeeding.

**Figure 3:**
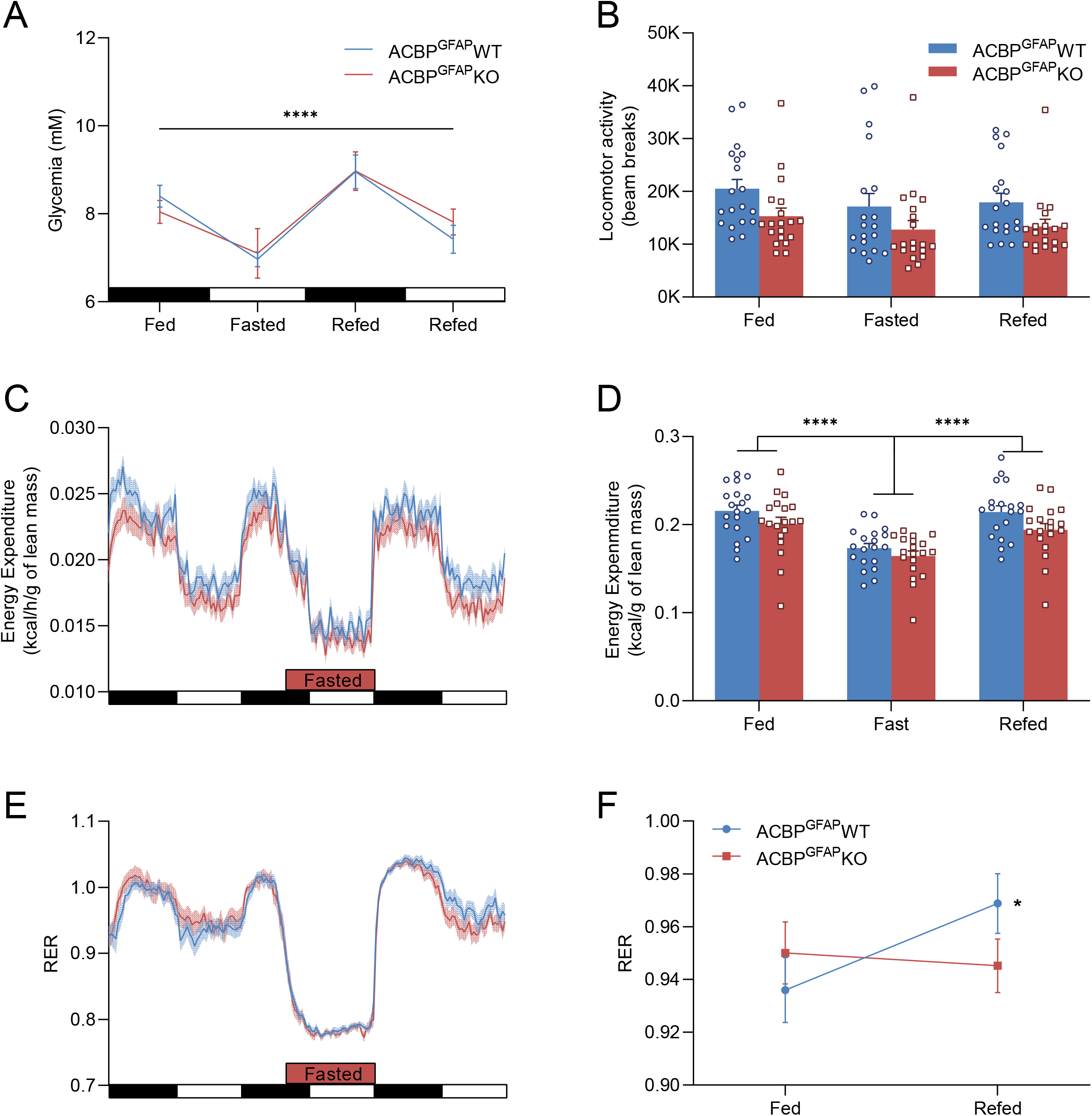
Energy metabolism flexibility in astroglial ACBP knockout mice subjected to fasting-refeeding. ACBP^GFAP^ WT and KO male mice were subjected to a 16h fast. (**A**) Glycaemia was measured 6h (ZT18) after the onset of dark cycle and 9h (ZT9) after onset of light cycle in both the fed and fasted states, N=8. (**B**) Locomotor activity, (**C-D**) Energy expenditure and (**E-F**) respiratory exchange ratio (RER) in ACBP^GFAP^ WT and KO mice in *ad libitum* fed, fasted and refed conditions. (**D**) Total energy expenditure in the different conditions and (**F**) Respiratory exchange ratio (RER) in the light cycle in fed and refed states, N=19. * p<0.05 (compared to fed WT), **** p<0.0001, Two-way ANOVA with Sidak post hoc test.

Although locomotor activity (Fig 3B) and energy expenditure (Fig 3C and D) trended lower in ACBP^GFAP^ KO mice in fed and refed states, it was not statistically significant. Fasting decreased energy expenditure (Fig 3C and D) and RER (Fig 3E and Suppl Fig1A) as expected, but these responses were similar in both groups. During refeeding, RER values are typically increased and can go above 1 due to increased carbohydrate utilization and lipogenesis. This response was observed in control mice when comparing RER values in the fed vs. refed state (Fig 3F). However RER did not increase to the same extent in the KO group to suggest decreased glucose utilization during refeeding (Fig 3F and Suppl Fig1A). Lower RER in the refeeding phase could result from the reduced food intake during refeeding in ACBP^GFAP^ KO mice (Fig 2D). Nevertheless, these data are in line with our previous finding that ICV administration of ODN increases glucose utilization in wild type male mice (19).

### Astroglial ACBP deletion alters feeding behavior during cold exposure

We and others previously reported that ODN directly increases the firing activity of POMC neurons (19, 23). In addition, ICV ODN promotes whole body glucose utilization and reduces feeding via the melanocortin pathway (19, 20). Given that metabolic responses to changes in housing temperature are partly dependent on the melanocortin pathway, we tested if male mice with astroglial ACBP deletion have impaired feeding and metabolic responses during cold exposure (4°C) or at thermoneutrality (30°C).

First, we assessed the impact of the housing temperature on hypothalamic ACBP expression in the ARC during both cold exposure (4°C) and thermoneutrality (30°C). Cold exposure during 24h lead to a significant increase in ACBP expression compared to normal temperature (21°C) and thermoneutrality (Fig 4A). AgRP expression was strongly induced by cold exposure while POMC expression was not affected by the temperature (Fig 4B).

**Figure 4:**
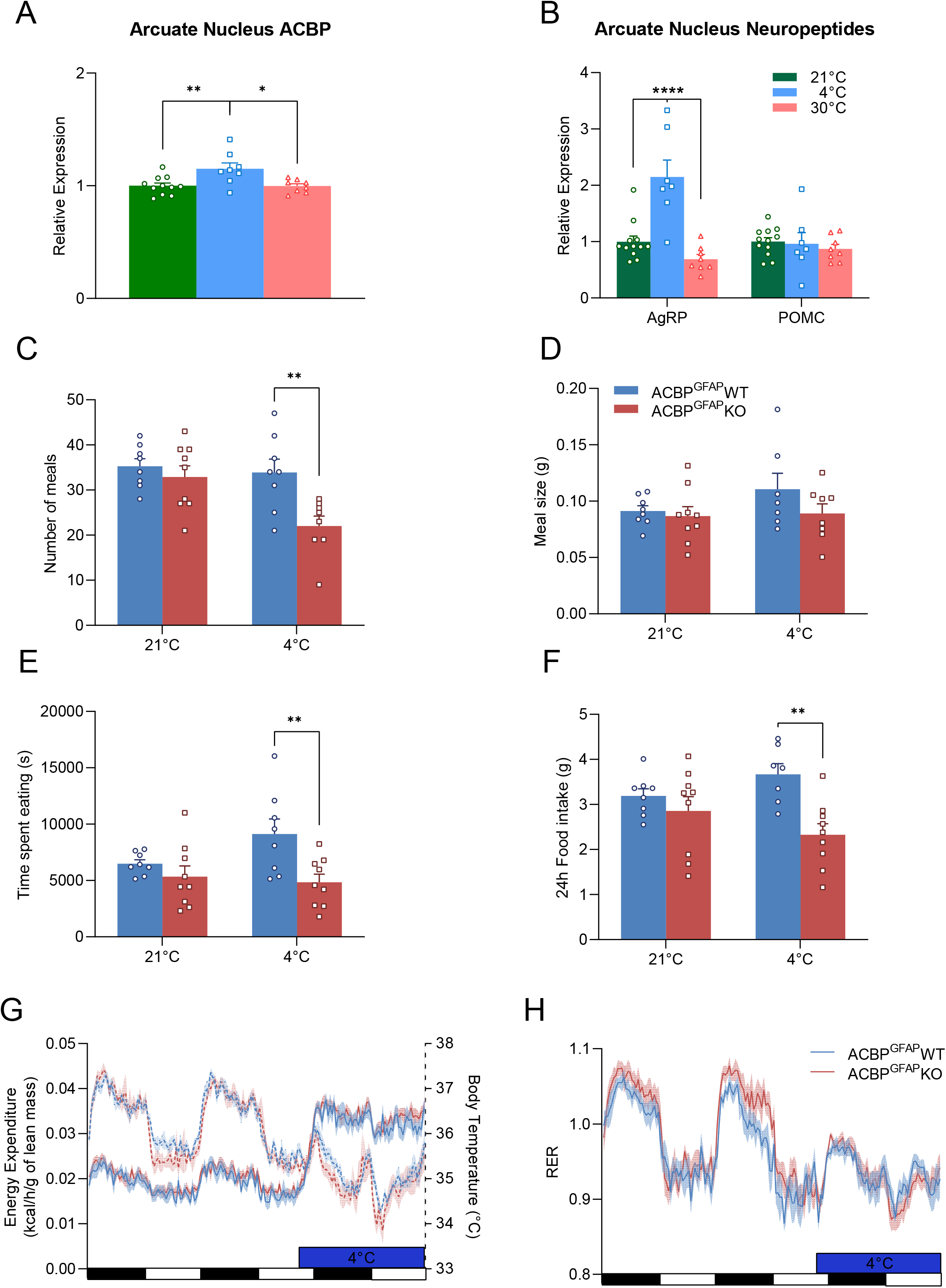
Feeding behavior and metabolic responses during cold exposure in astroglial ACBP knockout mice. (**A**) acbp, (**B**) agrp and pomc mRNA levels in the arcuate nucleus of male C57BL/6 mice exposed during 24h to normal housing temperature (21°C), cold (4°C) or thermoneutral conditions (30°C). * p<0.05, ** p<0.01 **** p<0.0001 One-(**A**) or Two-(**B**) Way ANOVA with Sidak post hoc test, N=8-12. ACBP^GFAP^ WT and KO male mice were housed at room temperature (21°C) and exposed to cold (4°C) in CLAMS metabolic cages. (**C**) Total number of meals, (**D**) Average meal size, (**E**) Total time spent eating and (**F**) Cumulative Food intake were measured during 24h at 21°C and 24h at 4°C. ** p<0.01, Two-way ANOVA with Sidak post hoc test compared to ACBP^GFAP^ WT (**G**) Energy expenditure, body temperature and (**H**) respiratory exchange ratio (RER) at 21°C and 4°C, N=8-9.

When assessing feeding behavior during cold exposure, we found that the total meal number (Fig 4C), time spent eating (Fig 4E) and cumulative food intake (Fig 4F) were significantly reduced in male ACBP^GFAP^ KO mice while the average meal size was not affected (Fig 4D). These findings therefore suggest that ACBP is required to mount an appropriate feeding response during energy-demanding conditions and are in line with the reduced food intake observed during the refeeding period (Fig 2D).

As expected, cold exposure (4°C) during 24h decreased body temperature and increased energy expenditure compared to regular housing temperature at 21°C (Fig 4G and Suppl Fig 1B-C). Cold exposure also led to a drop in RER induced by increased fatty acid utilization (Fig 4H and Suppl Fig 1D). However, these responses were not affected in ACBP^GFAP^ KO male mice compared to ACBP^GFAP^ WT.

A different cohort of mice housed in metabolic cages was transitioned from normal housing temperature (21 °C) to thermoneutrality (30°C). Thermoneutrality reduced all the feeding parameters analyzed (Fig 5A-D) in both genotypes leading to a decreased cumulative food intake which was more pronounced in ACBP^GFAP^ KO mice (Fig 5D). This response in ACBP^GFAP^ KO mice at thermoneutrality is related to a non-significant trend towards increased meal number and food intake in basal conditions (21°C), a pattern we have not observed in mouse cohorts in Fig 2 and 4. As expected, energy expenditure was decreased at thermoneutrality in control mice with a similar response in ACBP^GFAP^ KO mice (Fig 5E and Suppl Fig 1E). RER was also similar between groups at thermoneutrality (Fig 5F and Suppl Fig 1F).

**Figure 5:**
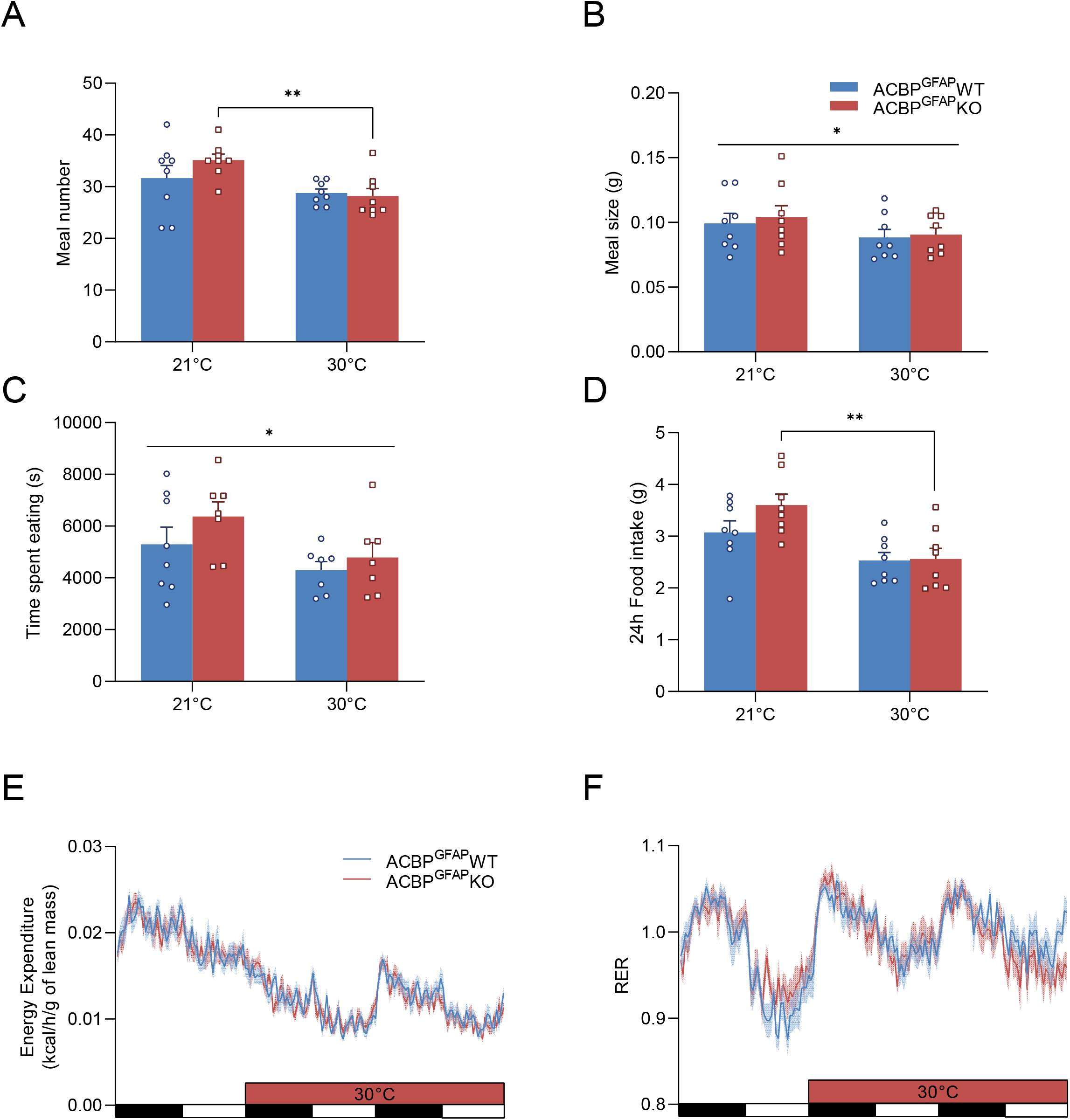
Feeding behavior and metabolic responses at thermoneutrality in astroglial ACBP knockout mice. ACBP^GFAP^ WT and KO male mice were housed at room temperature (21°C) and at thermoneutrality (30°C) in CLAMS metabolic cages. (**A**) Total number of meals, (**B**) Average meal size, (**C**) Total time spent eating and (**D**) Cumulative Food intake were measured during 24h at 21°C and 48h (averaged to 24h) at 30°C. (**E**) Energy expenditure and (**F**) respiratory exchange ratio (RER) at 21°C and 30°C. * p<0.05, Two-way ANOVA without or ** p<0.01 with Sidak post hoc test compared to 21°C, N=7-8.

## CONCLUSION

Despite a subtle change in carbohydrate utilization during refeeding (Fig 3F), astroglial ACBP deletion did not significantly impair adaptive metabolic responses (e.g. body temperature, energy expenditure) to acute challenges that are dependent on the melanocortin pathway. Importantly, our findings show that astroglial ACBP deletion impairs feeding responses to a fast and cold exposure, thereby suggesting that ACBP is required to promote feeding in response to acute challenges. These findings are in contrast with genetic and pharmacological studies, including ours, showing that ACBP and its derived peptide ODN reduce calorie intake (15). Indeed, our former (19) and current studies show that ACBP^GFAP^ KO mice have increased food intake during HFD compared to ACBP^GFAP^ WT controls (Fig 1A). It is however important to mention that it takes several weeks on HFD to observe a significant increase in cumulative calorie intake in ACBP^GFAP^ KO mice compared to control littermates. Accordingly, we showed that virally-mediated overexpression of ACBP in ARC GFAP astrocytes (~ 40% increase of ACBP expression) in adult wild type mice reduces cumulative food intake and body weight only ~ 7 to 8 weeks following viral injection in chow fed conditions (19). This suggests that genetic manipulation of ACBP expression in astrocytes affects cumulative calorie intake solely in the long term. These observations contrast with the rapid decrease in food intake induced by ICV ODN administration in mice and rats. Such differences between genetic and pharmacological models targeting ACBP are not surprising and could be explained by a different endozepinergic tone exerted by endogenous ACBP (weak) vs. exogenous ODN (strong) on POMC neurons and the melanocortin pathway and/or the recruitment of different neuronal populations. In addition, deletion or overexpression of ACBP in astrocytes is expected to affect both its role as a modulator of intracellular acyl-CoA metabolism (17) and as a secreted peptide making comparisons with the effect of ICV ODN difficult to interpret.

Nevertheless, our findings demonstrating that astroglial ACBP is required to mount the appropriate feeding response after a fast or during cold exposure were somewhat surprising given its reported POMC-dependent anorectic effect in different rodent models. However, the dogma that POMC neurons solely promote satiety has been recently discussed (25) and challenged by a study showing their activation can promote feeding. Indeed, endocannabinoids, that are well known to stimulate feeding, activate POMC neurons leading to increased food intake via the release of the hunger promoting POMC-derived beta-endorphin and stimulation of downstream mu-opioid receptor (26). Interestingly, fasting (27) and cold exposure (28) are known to increase circulating and/or hypothalamic endocannabinoid levels to stimulate feeding and regulate energy metabolism flexibility (29). It is thus tempting to speculate that in conditions associated with increased endocannabinoid tone such as cold and fasting, endogenous ACBP may potentiate the release of beta-endorphin by POMC neurons, rather than alpha-MSH, to promote feeding. Thus, the effect of endogenous ACBP on the type of POMC-derived neuropeptide secreted and feeding could be context dependent with hunger promoting properties in response to acute energy mobilizing challenges (e.g. fasting and cold) while it promotes satiety during chronic energy surfeit such as HFD (19, 23). The hunger-promoting effect of ACBP is in agreement with a recent study showing that circulating ACBP can stimulate feeding in mice (30).

Together, these results suggest that endogenous ACBP is not involved in metabolic flexibility but is required for feeding responses to acute challenges. These findings provide a new framework in which the role of astroglial ACBP on feeding behavior is dependent on the metabolic status.

## AUTHORS CONTRIBUTIONS

K.B., R.M., J. R. and D.R. helped with colony management, performed feeding and metabolic studies, and qPCR experiments. S.F. contributed to conceptualization, feeding results interpretation and contributed to manuscript preparation. K.B., R.M., J. R. and T.A. analyzed results and prepared the manuscript.

## ACKNOWLEDGEMENTS

We are grateful to Susanne Mandrup for the ACBP-floxed mice. We thank the CRCHUM rodent metabolic phenotyping core facility for their help with CLAMS and MRI studies. This work was supported by grants from the Canadian Institutes of Health Research (MOP115042 and PJT153035), Diabète Québec and Réseau de recherche en santé cardiométabolique, diabète & obésité from Fonds de Recherche Québec-Santé (CMDO-FRQS). T.A. was supported by a salary award from FRQS. R.M. and J. R. were respectively supported by doctoral and postdoctoral FRQS fellowships.

## DISCLOSURES

Authors have no conflict of interest to declare.

## DATA AVAILABILITY STATEMENT

Data are available on request from the authors

## METHODS

### Animals

Experimental animals were housed on a 12 h light – 12 h dark cycle (dark from 6:00 PM to 6:00 AM) in a pathogen-free environment. Housing temperature was maintained at 21°C (70°F) with *ad-libitum* access to water and standard irradiated chow diet. Regularly autoclaved cages were supplemented with nesting materials and cages were changed every two weeks. Colony health was monitored via a sentinel mouse exposed to feces from animals of the same rack.

Male C57BL/6J mice (000664) were purchased from Jackson Laboratory (8-10 weeks old). ACBP^+/+^;GFAP-cre (ACBP^GFAP^ WT) and ACBP^fl/fl^;GFAP-Cre (ACBP^GFAP^ KO) mice were generated as described previously (19). Experimental male mice (ACBP^GFAP^ WT and KO) were moved at 4 weeks of age to an experimental housing room on a reverse light-dark cycle (dark cycle from 10:00 AM to 10:00 PM). Mice were separated a few days before they were allocated to their experimental groups. For all studies, age-matched littermates were used and individually housed. Genotype, age, sex, and number of mice are indicated for each experiment in the corresponding figure legends or methods section. Upon completion of the studies, mice were anesthetized with ketamine/xylazine and blood was collected via cardiac puncture when necessary. Mice were then euthanized by decapitation prior to tissue collection. Only treatment naive mice were used at the time of study.

### High fat diet study

Five to six week old mice were individually housed and fed during 14 weeks with a high fat diet (HFD) (Modified AIN-93G purified rodent diet with 50 % kcal from fat derived from palm oil, Dyets, Bethlehem, PA, USA) and were switched to standard chow diet for an additional 4 weeks.

### Body composition analysis

Total fat and lean mass was assessed using a nuclear echo magnetic resonance imaging (EchoMRI) whole-body composition analyzer as described (19).

### Metabolic cages

Respiratory exchange ratio (RER), energy expenditure and locomotor activity were measured by indirect calorimetry in a Comprehensive Lab Animal Monitoring System metabolic cages (CLAMS, Columbus Instruments International, Columbus, OH, USA). Animals were single housed in CLAMS apparatus at 21±0.2°C in a dark-light cycle during 24 h to acclimate before any measurements. Following acclimation, energy balance parameters were measured during 48h. Energy expenditure was normalized by lean mass. For cold exposure studies, 10-12 week old naive mice were implanted with intra-peritoneal temperature probes and allowed to recover during 10-12 days before housing in CLAMS metabolic cages. Energy metabolism parameters were measured at 21°C during 24 h followed by 24 h at either 4°C, 24h or 48 h at 30°C. Mice were sacrificed and tissue collected in their respective temperature challenge conditions.

### Meal pattern analysis

Meal pattern analysis was performed on individual mouse data from .CSV files generated by the Oxymax software (Columbus Instruments International, Columbus, OH, USA) during housing in metabolic cages, compiled and analyzed with MATLAB (MathsWorks© R2021a, update3). Parameters for analysis were set in accordance with previous studies (31–33). A feeding bout was set at a minimum of 0.03 g of food consumed or lasting a minimum of 10 s. A meal was defined as a collection of bouts separated by less than 300 s intervals or inter-bout intervals.

### Real-time PCR

RNA extraction and real-time PCR was performed on fresh microdissections of the arcuate nucleus that include the median eminence as previously described (19) in male C57BL/6 mice exposed to different housing temperatures during 24h. Data were analyzed using the standard curve method and normalized to either 18s, actin and cyclophilin mRNA expression levels or their geometric means, as determined using NormFinder software (34).

### Statistical analysis

All statistical analysis were performed using the latest version of GraphPad Prism software. Intergroup comparisons were performed by One- or Two-Way ANOVA with Sidak post hoc tests as described in figure legends when appropriate. p < 0.05 was considered significant. All data are expressed as means ± SEM.

### Study approval

All procedures using animals were reviewed and approved by the institutional animal care and use committee (Comité Institutionnel de Protection de Animaux, protocol #CM20034TAs) of Centre de Recherche du Centre Hospitalier de l’Université de Montréal (CRCHUM).

---

**Supplemental Figure 1:**
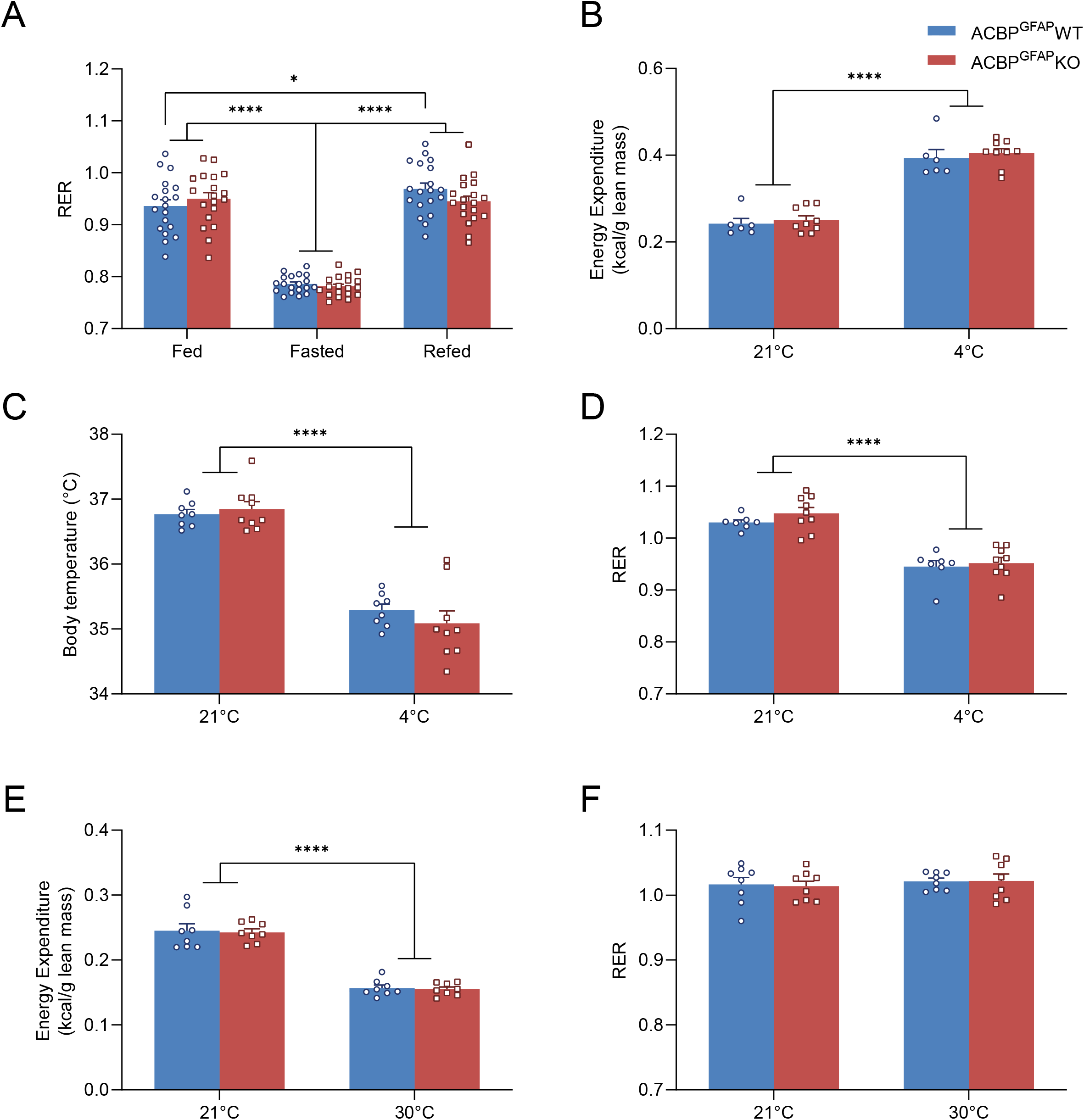
Energy metabolism parameters during challenges. **(A)** Average respiratory exchange ratio (RER) in ACBP^GFAP^ WT and KO male mice measured in the light cycle in fed, fasted and refed conditions, N=19. **(B)** Total energy expenditure, (**C)** body temperature and (**D)** average respiratory exchange ratio (RER) in ACBP^GFAP^ WT and KO male mice measured in the dark cycle at room temperature (21°C) and during cold exposure (4°C), N=6-9. **(E)** Total energy expenditure and (**F)** average RER in ACBP^GFAP^ WT and KO male mice measured in the dark cycle at room temperature (21°C) and thermoneutrality (30°C), N=8. * p<0.05 Two-Way ANOVA with Sidak post hoc test compared to WT fed state, **** p<0.0001 Two-Way ANOVA with Sidak post hoc test compared to fed state or 21°C.

## Notes

### Competing Interest Statement

The authors have declared no competing interest.

